# Synergism vs Additivity - Defining the Interactions between Common Disinfectants

**DOI:** 10.1101/2021.08.09.455772

**Authors:** Daniel J. Noel, C. William Keevil, Sandra A. Wilks

**Affiliations:** School for Biological Sciences, University of Southampton, Highfield Campus, Southampton, United Kingdom; School of Health Sciences, University of Southampton, Highfield Campus, Southampton, United Kingdom

## Abstract

Many of the most common disinfectant and sanitizer products are formulations of multiple antimicrobial compounds. Products claiming to contain synergistic formulations are common, although there is often little supporting evidence. The antimicrobial interactions of all pairwise combinations of common disinfectants (benzalkonium chloride, didecyldimethylammonium chloride, polyhexamethylene biguanide, chlorocresol and bronopol) were classified via checkerboard and validated by time-kill analyses. Combinations were tested against *Acinetobacter baumannii* NCTC 12156, *Enterococcus faecalis* NCTC 13379, *Klebsiella pneumoniae* NCTC 13443 and *Staphylococcus aureus* NCTC 13143. Synergistic interactions were only identified between cholorocresol with benzalkonium chloride, and chlorocresol with polyhexamethylene biguanide. Synergism was not ubiquitously demonstrated against all species tested and was on the borderline of the synergism threshold. These data demonstrate that synergism between disinfectants is uncommon and circumstantial. Most of the antimicrobial interactions tested were characterised as additive. We suggest that this is due to the broad, non-specific mechanisms associated with disinfectants not providing opportunity for the combined activities of these compounds to exceed the sum of their parts.

**IMPORTANCE:** The scarcity of observed synergistic interactions suggests that many disinfectant-based products may be misinterpreting combined mechanisms of interaction. We emphasise the need to correctly differentiate between additivity and synergism in antimicrobial formulations, as inappropriate classification may lead to unnecessary issues in the event of regulatory changes. Furthermore, we question the need to focus on synergism and disregard additivity when considering combinations of disinfectants, as the benefits that synergistic interactions provide are not necessarily relevant to the application of the final product.

## INTRODUCTION

Under the US Federal Insecticide, Fungicide, and Rodenticide Act (FIFRA), an antimicrobial pesticide is defined as a biocide that disinfects, sanitizes, reduces, or mitigates growth or development of microorganisms (1). Depending on the application, any antimicrobial pesticides to be sold or distributed in the USA must be registered with either the Environmental Protection Agency (EPA) or Food and Drug Administration (FDA) to ensure the product meets minimum efficacy and safety standards. Equivalent legislation can be found globally, for example the European Union enforces the Biocidal Products Regulations (BPR) (2). Inadvertently, the implementation of pesticide regulations have effectively stopped most research into novel antimicrobial compounds due to the unfavourable cost of development and registration (3). It remains more financially viable for companies to develop formulations containing currently approved active compounds than to risk the cost of developing and attempting to authorise novel antimicrobials. In the EU this remains the case even after the BPR revision in 2012 which aimed, among other things, to simplify the process of product authorisation (2).

As a result, many of the most widely available antimicrobial disinfectant and sanitizer products consist of combinations of a limited number of individual compounds. These products are routinely used as disinfectants and antiseptics in healthcare settings, industrial environments and in day-to-day life in the form of surface sprays, wipes and hand sanitizers. The central axiom that synergistic interactions can occur between antimicrobials with different mechanisms of action and target sites has resulted in the liberal use of claims of synergy when describing and marketing multi-component disinfectants products.

However, correctly classifying the type of interaction between antimicrobial agents is a challenging process. For example, inconsistencies regarding the classification of an antimicrobial interaction can arise depending on the method employed (4–7). Common techniques used to investigate antimicrobial interactions include the E test, time kill and checkerboard methods.

Of these methods, the most widely used is the checkerboard assay; a variation of the broth microdilution technique to determine the minimum inhibitory concentration (MIC). In brief, each compound is serially diluted along either the X or the Y axis of a multi-well plate containing growth medium. The wells are then inoculated with the test species and incubated. The output of the test is the fractional inhibitory concentration index (FICI). The lower the FICI value, the higher the level of interaction between the two tested compounds.

The checkerboard method provides a high-throughput technique that can yield a large amount of information about how pairs of antimicrobials interact in a relatively short period of time. Despite this the checkerboard method does raise significant challenges when it comes to interpreting results and thus classifying antimicrobial interactions. Firstly, the outcome of a checkerboard assay varies depending on the method of interpretation (8), which is especially significant when many published articles do not explicitly state the method of interpretation used. In addition, the FICI thresholds that can be set to define the verdict of an interaction often vary between publications, creating issues around standardisation and comparability of results.

Further complications arise when comparing data between species or strains, with publications reporting significant variations in the classification of combined activities of mixtures of both disinfectants (9, 10) and antibiotics (11, 12). This has resulted in the same combinations being reported as synergistic, additive or indifferent depending on the species or even strain it was tested upon. While certain academic journals have implemented FICI standards when reporting checkerboard data (13–15), these are not universally adhered to between journals which further contributes to inconsistencies between publications.

A lack of universal consensus on the definitions of antimicrobial interactions provides additional issues. For the purposes of this publication, the definitions used are in accordance with those set out by the European Committee on Antimicrobial Susceptibility Testing (EUCAST). According to EUCAST, an indifferent interaction is one whereby the activity of both components combined is equal to the activity of the most active component (16). An additive interaction has a combined activity of no greater than the sum of the activities of each component, while the sum of the individual activities has to be exceeded by the combined activity in order for the interaction to be classed as synergistic (16). Antagonism is the inverse, whereby the activity of both components combined is lower than the most active component (16).

These definitions are not universally accepted or adhered to, for example multiple leading journals in the field do not accept ‘additive’ checkerboard interpretations due to the definition being commonly misunderstood and the intrinsic variability of the method (13, 15). In the guidance to authors it is even suggested that alternative terms are used such as “nonsynergistic” (15), thus encouraging researchers to disregard intermediate levels of activity and focus exclusively on interactions that demonstrate synergy (15, 14). This does not mean that these journals completely disregard the existence of additive interactions, instead that additivity is too difficult to pinpoint and identify reliably using the checkerboard methodology.

The confusion between additivity and synergism and the methods employed to distinguish between them has led to the two terms often being used interchangeably and potentially erroneously. The confusion is especially significant as synergism in an antimicrobial formulation is considered “surprising” and thus is patentable (3). This, alongside the increased marketability the “synergy” buzzword brings, provides a commercial incentive to classify such formulations as synergistic, even if the evidence is circumstantial and the definitions are misunderstood. In addition, results that support patentable ideas often remain unpublished in order to prevent potential loss of intellectual property. This means that the evidence required to support a patent is not often subjected to the same scrutiny as peer-reviewed publications.

These factors combined lead to academic and commercial-related research being “all or nothing” - exclusively focusing on synergistic antimicrobial interactions and completely disregarding additive interactions.

With these issues in mind, this study classifies the nature of the interactions between antimicrobials that are commonly used in disinfectant and sanitizer formulations. The compounds examined in this study are listed in Table 1, alongside their mechanisms of action, applications and compounds they are commonly found with in formulations. Previous research has indicated variability between species and strains (9–12), therefore clinically-relevant bacterial species that display a degree of antibiotic resistance were selected in order to provide a stringent test. In addition, strict activity classification thresholds were used to provide clarity and to maintain consistency with the standards set by leading journals in the field (13–15). Additivity was included as a classification due to the context of the test.

**Table 1.** Summary of characteristics of the disinfectants used in this study.

| Compound | Cellular target | Antimicrobial mechanism | Applications | Compounds commonly associated with in formulations |
| --- | --- | --- | --- | --- |
| Benzalkonium chloride (BAC) | Membrane | Positively charged quaternary nitrogen interact with anionic lipids, facilitating its own uptake. Adsorption allows hydrophobic tails to insert into the bilayer, causing disruption of lipid organisation and breaches in the permeability barrier. Leads to leakage of low molecular weight material, loss of proton motive force and uncoupling of oxidative phosphorylation. (21, 22, 37–39) | Surface disinfection sprays and wipes, eye/ear drops, burn treatments. | DDAC, PHMB, ethanol. |
| Didecyl-dimethyl-ammonium chloride (DDAC) | Membrane | Positively charged quaternary nitrogen interact with anionic lipids, facilitating its own uptake. Adsorption allows hydrophobic tails to insert into the bilayer, causing disruption of lipid organisation and breaches in the permeability barrier. Leads to leakage of low molecular weight material, loss of proton motive force and uncoupling of oxidative phosphorylation. (21, 37, 39, 40) | Surface disinfection sprays and wipes, sterilization of surgical equipment. | BAC, PHMB, ethanol. |
| Polyhexamethylene biguanide (PHMB) | Membrane | Biguanide group interacts and sequesters anionic lipids, forming homogenous lipid domains. This disrupts bilayer organisation and leads to permeability of membrane and intracellular leakage (22, 39, 41–43). Evidence also suggests that PHMB is able to translocate across the bacterial membrane and condense bacterial DNA and prevent DNA replication (44, 45). | Surface disinfection sprays and wipes, wound dressings, contact lens cleaning solution, swimming pool cleaners. | BAC, DDAC, ethanol. |
| Bronopol | Proteins.<br>ROS generated target macromolecular structures | Catalyses oxidation of thiols to di-sulphides, cross-linking proteins. Changes to protein structure result in impeded functionality. This reaction also produces reactive oxygen species which damage intracellular structures. (46) | Disinfectant, preservative. | BAC, DDAC. |
| Chlorocresol | Membrane | Disruption of the permeable barrier and induce leakage of low molecular weight intracellular components. Leads to loss of proton motive force and uncoupling of oxidative phosphorylation. (21, 47) | Antiseptic, preservative. | Ethanol, triclosan. |

## RESULTS

### Minimum inhibitory concentration

The MIC results are summarised in Table 2. Benzalkonium chloride (BAC), didecyldimethylammonium chloride (DDAC), polyhexamethylene biguanide (PHMB) and bronopol achieved MIC values in the range of 31 μg/ml to 2 μg/ml across all bacterial species tested, while chlorocresol achieved MIC values in the considerably higher range of 600 μg/ml to 125 μg/ml. DDAC achieved an MIC that ranged from 8 μg/ml to 2 μg/ml across all tested species. BAC achieved MIC values of 8 μg/ml and 5 μg/ml for *E. faecalis* and *S. aureus* respectively, while the MIC values for *A. baumanii* and *K. pneumoniae* were significantly higher at 31 μg/ml and 20 μg/ml respectively. Chlorocresol achieved MIC values of 200 μg/ml and 125 μg/ml for *K. pneumoniae* and *A. baumanii* respectively, while the MIC values for *S. aureus* and *E. faecalis* were significantly higher at 600 μg/ml and 500 μg/ml respectively. Bronopol followed a similar trend with MIC values of 8 μg/ml and 4 μg/ml for *K. pneumoniae* and *A. baumanii* respectively and 20 μg/ml and 16 μg/ml for *S. aureus* and *E. faecalis* respectively.

**Table 2.** Minimum inhibitory concentration values of common disinfectants against clinically relevant bacterial species.

| Bacterial species | MIC (µg/ml) |  |  |  |  |
| --- | --- | --- | --- | --- | --- |
|  | Benzalkonium chloride | Didecyldimethyl-ammonium chloride | Polyhexamethylene biguanide hydrochloride | Bronopol | Chlorocresol |
| *A. baumannii* NCTC 12156 | 31 | 8 | 16 | 4 | 125 |
| *E. faecalis* NCTC 13379 | 8 | 4 | 8 | 16 | 500 |
| *K. pneumoniae* NCTC 13443 | 20 | 6 | 6 | 8 | 200 |
| *S. aureus* NCTC 13143 | 4 | 2 | 6 | 20 | 600 |

The MIC for DMSO was greater than 100,000 μg/ml (10 % v/v) for all species tested (results not shown). DMSO was therefore not responsible for the activity demonstrated by chlorocresol.

### Checkerboard assay

The classification of interactions between pairs of disinfectants were evaluated via the checkerboard method. Results are summarised in Table 3 and Figure 1. The combination of BAC + chlorocresol demonstrated synergism against *S. aureus* and *E. faecalis*, and PHMB + chlorocresol in combination proved synergistic against *E. faecalis*. The FICI values of these interactions were all on the threshold of the “synergistic” classification (0.5), therefore this synergism was considered borderline.

**Table 3.**
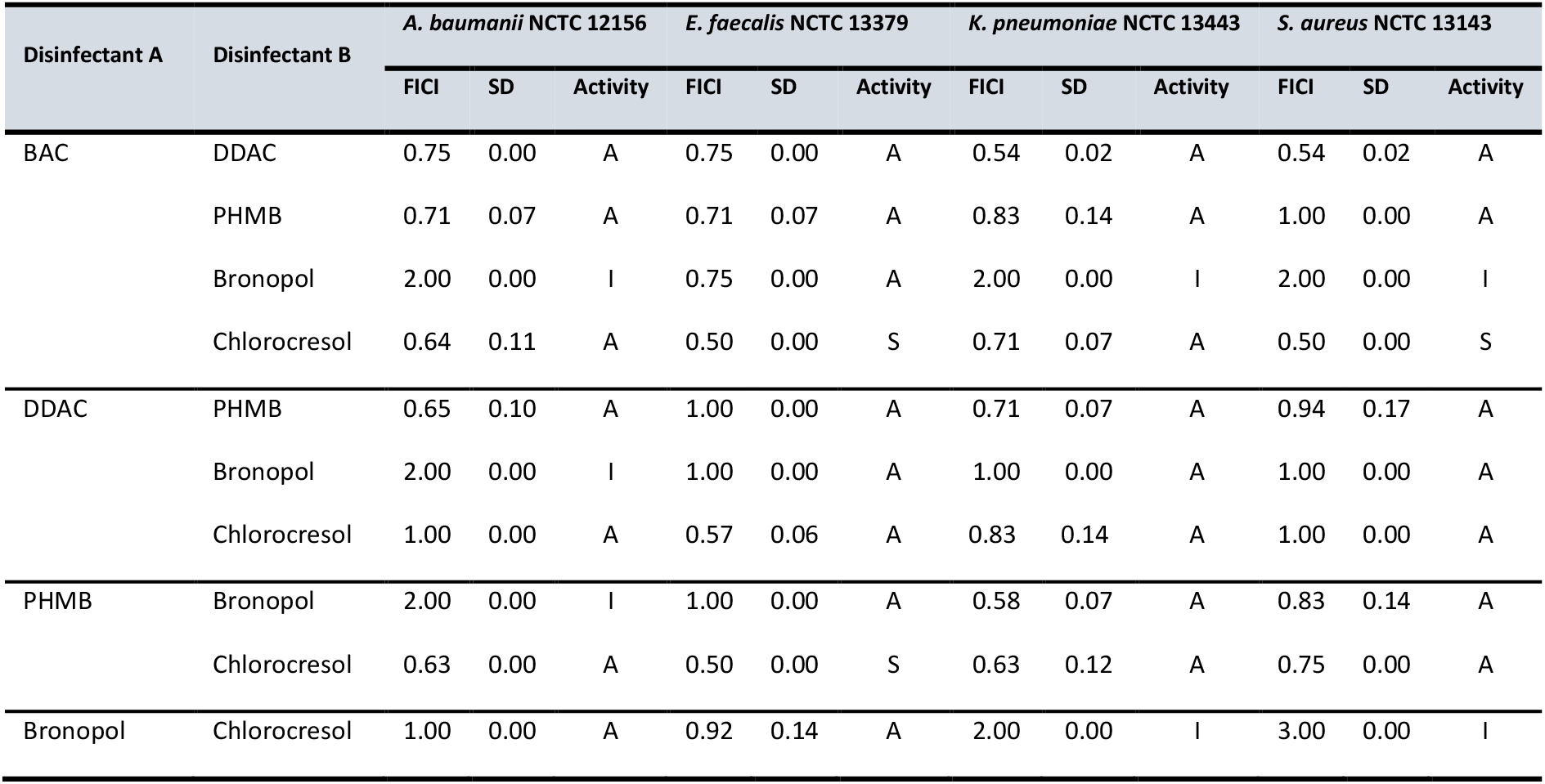
Combined antimicrobial activities of pairwise combinations of five common disinfectants.^a^ ^a^Abbreviations: FICI – Fractional Inhibitory Concentration Index. SD – standard deviation. BAC – benzalkonium chloride. DDAC - didecyldimethylammonium chloride. PHMB - polyhexamethylene biguanide hydrochloride. A – additive. I – indifferent. S – synergistic.

**Figure 1.**
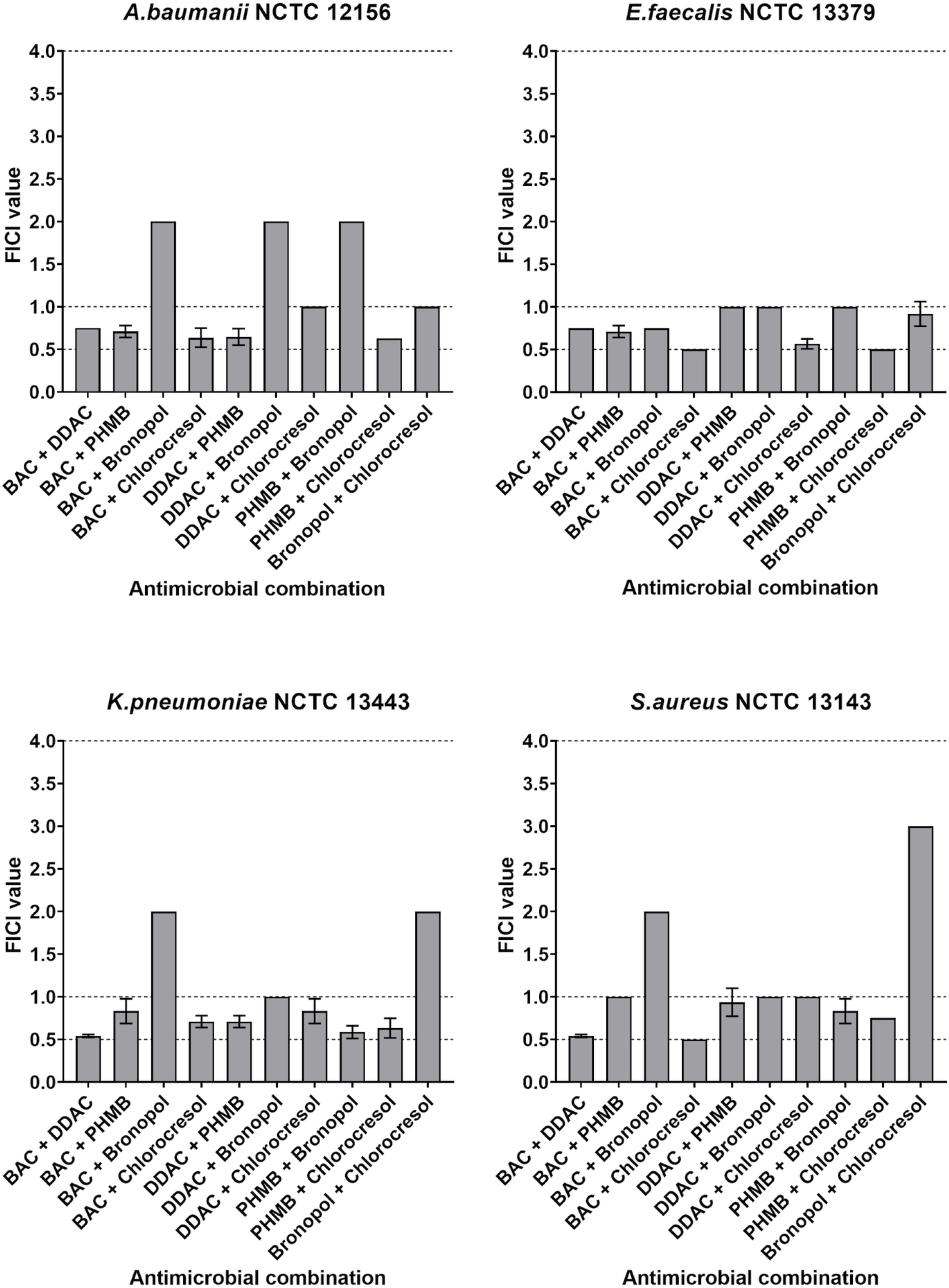
Fractional inhibitory concentration Indices (FICIs) of combinations of five common antimicrobial disinfectants. Dotted lines depict the thresholds between synergism (FICI ≤ 0.5), additivity (0.5 < FICI ≤ 1.0) and indifference (1.0 < FICI ≤ 4.0).

All combinations of antimicrobials tested demonstrated varying degrees of additivity against at least one of the tested species. Most notably, every combination of cationic membrane-active antimicrobials (BAC, DDAC and PHMB) demonstrated an additive mechanism across all species tested.

Any disinfectant combination that included bronopol produced combined activities that were inconsistent across the different species, with all combinations demonstrating both indifference and additivity that varied in an inconsistent species-dependant manner. Antimicrobial combinations including bronopol were responsible for every indifferent combination tested.

### Time-Kill Assay

Disinfectant combinations previously identified as synergistic were analysed via the time-kill methodology. All three combinations demonstrated a ≥2 log^10^ reduction in CFU/ml compared to the most active constituent alone and the initial inoculum after 24 hours, thus validating the synergistic interactions demonstrated by these three combinations (Figure 2).

**Figure 2.**
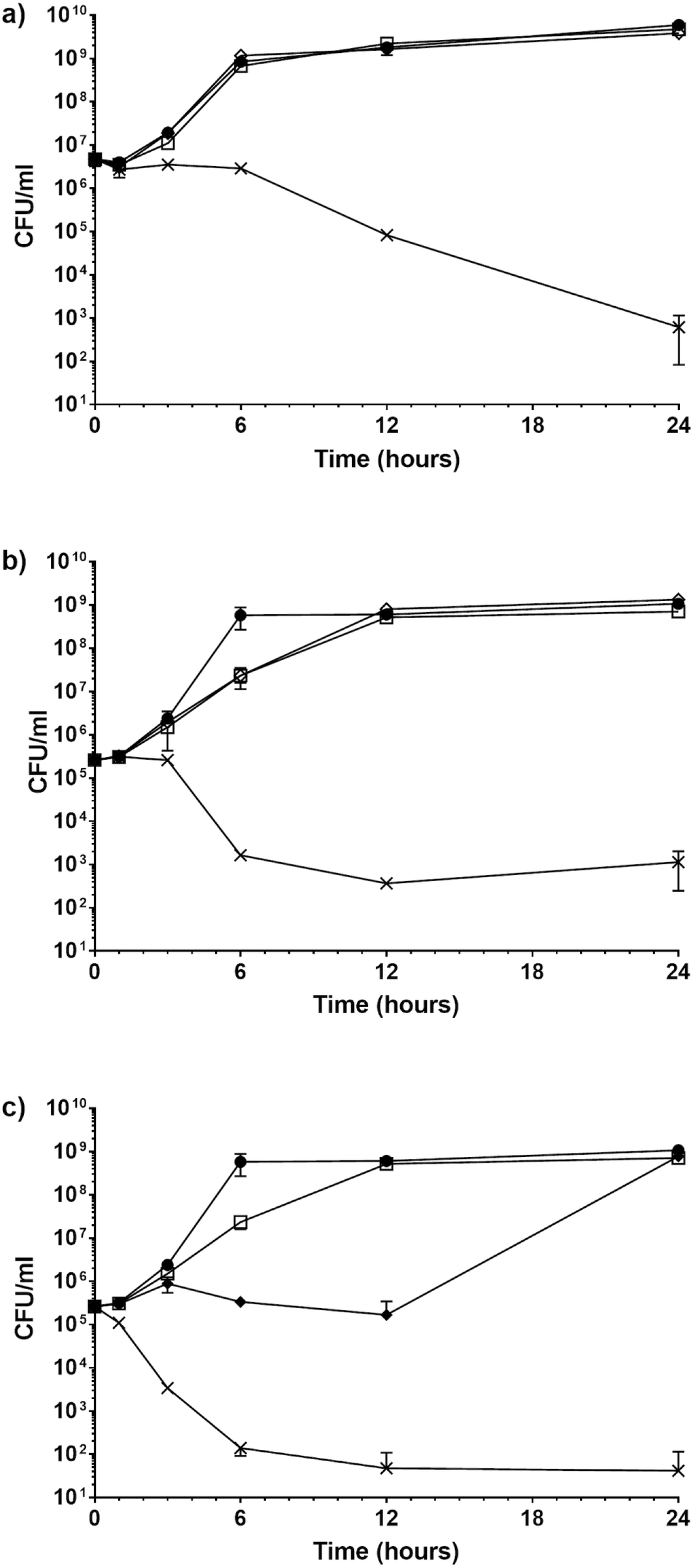
Time-kill curves of synergistic combinations of disinfectants. a) *Staphylococcus aureus* NCTC: 13143 exposed to a combination of benzalkonium chloride and chlorocresol. •, growth control; ⬦, 0.0001% v/v benzalkonium chloride; □, 0.01% v/v chlorocresol; X, 0.0001% v/v benzalkonium chloride and 0.01% v/v chlorocresol in combination. b) *Enterococcus faecalis* NCTC: 13379 exposed to a combination of benzalkonium chloride and chlorocresol. •, growth control; ⬦, 0.0002% v/v benzalkonium chloride; □, 0.0125% v/v chlorocresol; X, 0.0002% v/v benzalkonium chloride and 0.0125% v/v chlorocresol in combination. c) *Enterococcus faecalis* NCTC: 13379 exposed to a combination of polyhexamethylene biguanide and chlorocresol. •, growth control; ⬥, 0.0002% v/v polyhexamthylene biguanide; □, 0.0125% v/v chlorocresol; X, 0.0002% v/v polyhexamthylene biguanide and 0.0125% v/v chlorocresol in combination. All experiments were performed in triplicate.

*E. faecalis* displayed an impeded rate of growth when exposed to the individual disinfectants (Figure 2b, c), even though they were each below their respective MICs. The cultures demonstrated impeded growth up to 12 hours after exposure in comparison to the growth controls (Figure 2b, c). Despite this, the outcome at the 24-hour time point was sufficient for the combined activities of the disinfectants to each demonstrate a synergistic interaction.

## DISCUSSION

Many antimicrobial products used in medical, industrial and domestic environments consist of formulations of multiple individual disinfectants. Claims are often made regarding the synergistic mechanism of such formulations based on the various compounds present in the solution demonstrating varying mechanisms of action.

Despite these claims there is limited evidence to support the synergistic interactions between many of the most common disinfectants. In addition, previous reports indicate that the various methods employed to investigate these interactions can produce inconsistent results (4–7), and varying thresholds are often implemented to distinguish between synergistic, additive or indifferent mechanisms which can ultimately lead to variation between publications (17–19).

For example Soudeiha *et al*. reports an ‘additive’ interaction between colistin and meropenem when tested against *A. baumannii* clinical isolates in vitro, with ‘additive’ FICI values ranging from 0.61 to 1.83 (17). In contrast, when the same antimicrobial combination was tested against *A. baumannii* clinical isolates by Oliva *et al*., many of the interactions were reported as ‘indifferent’, despite the FICI values often being lower than those reported by Soudeiha *et al*. (17, 18). These discrepancies in the classification of antimicrobial interactions are simply due to different thresholds being used.

In addition, neither of the studies explicitly define the terms used (synergy, additive or indifference) to classify the antimicrobial interaction between colistin and meropenem (17, 18). Another similar study on the same antimicrobials conducted by Kheshti *et al*. reported FICI values of between 0.5 and 1 as “partial synergism” (19). The lack of clarification introduces additional ambiguity and hinders the ability to draw an overall conclusion between the published reports (17–19). While these examples investigate antibiotics specifically, the underlying issues extend to all antimicrobial interactions that are examined via the checkerboard methodology.

Collectively, these factors could lead to the incorrect classification of a combined antimicrobial activity. This is significant as it may result in consumers placing too much faith in a product, leading to potential inappropriate and ineffective usage. The goal of this study was to investigate the type of interaction between common disinfectants present in formulations, and highlight current issues surrounding antimicrobial interaction testing.

The nature of the interactions between 5 common disinfectants when used in pairwise combinations were classified via the widely used checkerboard method. The characteristics of the antimicrobial compounds used in this study are summarised in Table 1. A synergistic interaction between BAC and chlorocresol was observed against *E. faecalis* and *S. aureus*, and between PHMB and chlorocresol against *E. faecalis* (Table 3 and Figure 1). These combinations did not demonstrate synergism against *A. baumanii* or *K. pneumoniae* however, indicating that this synergistic mechanism is species-specific.

These synergistic combinations were tested further via the time-kill methodology. The antimicrobials combined resulted in a ≥5 log reduction in CFU/ml after 24 hours compared to when they were used individually, confirming synergistic interactions (Figure 2).

Indifference was observed in various combinations that contained bronopol, although it is important to note that these indifferent interactions were not consistent across the species tested (Table 3 and Figure 1). It has previously been reported that BAC and bronopol synergistically inhibit sulfide production in sulfate-reducing bacteria (20). As antimicrobial activity was measured via sulfide production it is difficult to draw comparisons between the results. This variation between reports further demonstrates that disinfectant interactions are not ubiquitous and vary in nature depending on the test species and methodologies used.

Interestingly, it was observed that every combination of cationic membrane-active antimicrobial (BAC, DDAC, PHMB) interacted additively across all species tested, with FICI values ranging consistently between 0.54 and 1.00. This suggests that disinfectants with similar mechanisms and cellular targets (see Table 1) consistently benefit from being in combination, although not to the point of synergism. We propose that this is due to similar-acting compounds having a limited but consistent scope to complement each other’s activities when used in combination. At the sub-lethal concentrations tested, the cationic membrane-active compounds both disrupt membrane stability and cause intracellular leakage (21–23) thus will each mechanistically benefit from the presence of the other. With broadly overlapping mechanisms the combined activity never has the opportunity to be greater than the sum of its parts, therefore the interaction is limited to additivity.

Of the 40 test conditions tested, 30 demonstrated an additive interaction against the respective target species (Table 3). In addition, all disinfectant combinations demonstrated at least one additive interaction against the various species. We believe that the abundance of additive antimicrobial interactions is due to the broad, non-selective mechanisms demonstrated by disinfectants. The wide range of cellular targets and high level of activity leaves little room for other additional disinfectant compounds to provide a suitably varying mechanism that would enable a synergistic interaction. As a result, any combined activities would simply be accumulative and would rarely ever be greater than the sum of its parts. Thus the majority of the interactions observed are additive. We therefore postulate that finding synergistic combinations of antimicrobials is more challenging when investigating compounds that have non-specific, broad mechanisms (for example disinfectants) than those that have more specific mechanisms of action (antibiotics).

Despite this observed scarcity in synergistic interactions, claims of synergy are common in disinfectant products. It is possible that products may be being inappropriately classified as synergistic due to the commercial incentives surrounding a “synergistic” claim combined with non-peer-reviewed supporting data and a lack of understanding of the terminology. Clarifying and identifying the differences between any additive and synergistic mechanisms within a disinfectant formulation is of vital importance and should not be dismissed. Synergistic combinations may provide unique and powerful activities that influence not only the effectiveness of the formulation, but impact how it can be effectively used. Overstating the effectiveness of such formulations by erroneously identifying interactions as ‘synergistic’ can lead to consumers placing too much faith in a product, which could lead to inappropriate use.

Furthermore, disinfectants are often under scrutiny by regulatory bodies and can be tightly controlled or withdrawn from use. Regulations vary between countries and regions, and are often reviewed and changed, for example in 2016 and 2017 the FDA banned a total of 24 active ingredients including triclosan for use in soaps (24, 25). Two of these listed active ingredients applied to specific antimicrobial combinations (25). Other active compounds, including benzalkonium chloride, have had their FDA rulings deferred on a year-by-year basis since 2016 at the request of manufacturers (26–31). This is in order to complete ongoing research into the safety and effectiveness, and as of time of writing the most recent deferral will expire on October 31^st^ 2021 (30).

The uncertainty surrounding biocide regulations creates a need for international products to be able to adapt and change their formulations to conform to local regulations. Making a substitution in a formulation is incredibly challenging if the component relies upon a synergistic interaction, as our data suggests that such interactions are very specific and uncommon (Table 3). In contrast, substituting an antimicrobial that provided an additive interaction to a mixture can be achieved relatively easily via a functional analogue, as such interactions are relatively common (Table 3). A formulation that has been inappropriately characterised as synergistic could lead to unnecessary challenge and expense if legislations change and a key component needs substituting. For these reasons fully understanding the nature of antimicrobial interactions is of paramount importance both to the commercial sector and to consumers.

The observed scarcity of synergistic interactions between broad-spectrum disinfectants also raises the question of whether the benefits of synergistic interactions outweigh the challenges required to identify them. In short, is it worth it? The obvious answer is yes, as there are significant benefits of synergistic interactions. Perhaps most obviously is enhanced antimicrobial activity leading to a higher efficacy, meaning a more effective and reliable product. However, broad-spectrum antimicrobial formulations contain concentrations of active compounds that are typically multiple orders of magnitude greater than the MIC of any likely target organism, and therefore efficacy is not usually a limitation that needs addressing. For example, most supermarket-branded antibacterial sprays and wipes contain between 1,000-20,000ppm BAC, while the MICs against clinically relevant bacterial species lie multiple orders of magnitude lower in the ranges of 4-31 ppm (Table 2). Additionally, there are many widely used disinfectants available that only contain one active component, therefore demonstrating that combined antimicrobial interactions are not *necessary* for a product to be effective and successful.

A second advantage of synergistic interactions is the ability to minimise resistance development, as targets would have to become resistant to multiple distinct mechanisms simultaneously (11, 32, 33). However, this benefit is not unique to synergistic interactions – it also applies to additive and even indifferent interactions too. Furthermore, resistance to disinfectants at optimal concentrations is not a widespread issue that regularly impacts the efficacy of products; thus it could be argued that this advantage is (currently) a moot point.

An additional advantage of a unique synergistic interaction is that it could enable compounds to be effective against entirely new targets that they otherwise could not. However, to our knowledge there is little evidence to demonstrate this occurring when considering disinfectant combinations specifically.

With multiple academic journals not accepting additive interactions (13) the current bar is set at distinguishing between synergistic interactions and everything else. However, synergistic interactions do not necessarily provide any discernible advantages over additive interactions when considering the quality and functionality of a disinfectant product. We therefore question whether these standards are necessary and suggest that the focus instead be shifted to distinguishing between additive and indifferent interactions when assessing the combined activity of broad-spectrum disinfectants.

It is important to note that disinfectant products routinely contain more than two ‘active’ components, alongside ‘inactive’ additives such as solvents, surfactants, emulsion stabilisers and fragrance enhancers. Such formulations therefore contain a network of complex interactions between multiple ‘active’ and ‘inactive’ compounds which will inevitably influence the overall product efficacy. To our knowledge the interactions between common ‘inactive’ components and ‘active’ disinfectants within a formulation have not been explored in the literature. Furthermore, the characterisation of complex interaction networks between combinations of more than two disinfectants has not been investigated. This study comprehensively and systematically classifies the interactions between common disinfectants, representing an important initial step toward fully elucidating the interaction networks that underpin the efficacy of disinfectant formulations used ubiquitously across the world.

## CONCLUSION

Disinfectant formulations are globally depended upon in healthcare environments, industrial settings and in day to day life. Their use as an infection control measure is critical, especially as the world looks for sustainable routes out of the COVID-19 pandemic. Many common formulations claim to be or are advertised as synergistic. However, the definitions surrounding synergism, additivity and indifference between antimicrobial compounds are often poorly understood and regularly misused. Understanding and not overstating the nature of these interactions is critical because it influences the correct usage of antimicrobial formulations, and also dictates the viability of substituting antimicrobials for functional analogues in the event of regulatory changes.

These data demonstrate that synergism between common disinfectants is a rare occurrence and any synergistic mechanisms are not necessarily ubiquitous across bacterial species. The majority of the interactions were characterised as additive, which we suggest is likely due to the broad range of cellular targets providing little opportunity for any given antimicrobial combination to be greater than the sum of its parts. We therefore question whether the current emphasis on synergistic interactions in academia and product development is necessary in the context of broad-spectrum disinfectants. Synergistic interactions are not likely to provide any discernible or impactful benefit over additive interactions in terms of the quality of the final product.

## MATERIALS AND METHODS

### Bacterial strains and growth media

The following bacterial strains were used in this study: *Acinetobacter baumannii* NCTC 12156, *Enterococcus faecalis* NCTC 13379, *Klebsiella pneumoniae* NCTC 13443 and *Staphylococcus aureus* NCTC 13143. The strains were selected due to their clinical relevance and impact on healthcare-associated infections (34). All bacterial strains were cultured in 10 ml Mueller-Hinton Broth (MHB) (Thermo Scientific) overnight at 37°C. Bacterial stocks were standardised to a final test suspension of 5 × 10^5^ CFU/mL.

### Stock solutions of antimicrobial compounds

Antimicrobial compounds were selected based on their presence in commercial antimicrobial formulations. BAC and DDAC are both quaternary ammonium compounds commonly found as components in antimicrobial formulations. PHMB and phenol derivatives such as chlorocresol are also common components. Bronopol was selected as it acts via a different mechanism in comparison to the other selected compounds. The characteristics of these antimicrobial compounds are summarised in Table 1.

Benzalkonium chloride (Thor Specialities Limited), didecyldimethylammonium chloride (Thor Specialities Limited), polyhexamethylene biguanide (Thor Specialities Limited) and bronopol (Thor Specialities Limited) were made up to a stock concentration of 10,000 μg/ml in ddH_2_O immediately before testing. Chlorocresol (Lanxess Limited) was made up to a stock concentration of 10,000 μg/ml in undiluted DMSO (Corning) immediately before testing.

### Minimum inhibitory concentration

The MIC values were determined using the broth microdilution method as described by the Clinical Laboratory Standards Institute (CLSI) (35). Due to the antimicrobial compounds demonstrating a wide range of potential activities, serial dilutions began from 10,000 μg/ml instead of 128 μg/ml as recommended for antibiotics. Experimentation was performed using 96-well plates in triplicate. Plates were incubated at 37°C overnight. The MIC was defined as the lowest concentration of active compound that completely inhibited bacterial growth in the microdilution wells as detected by the unaided eye when the bacterial growth in blank wells was sufficient. The MIC of DMSO was calculated for all tested species to ensure validity of chlorocresol MICs.

### Checkerboard assay

The checkerboard assay was used to determine the activities of antimicrobial compounds in combination as described previously (5, 8). Each well of a 96-well plate contained a final volume of 200 μl. Arrangements of antimicrobial compounds were made whereby one compound was serially-diluted twofold on the horizontal axis and another on the vertical axis, with final concentrations ranging from 4 x to 1/128 x MIC. Once prepared, each checkerboard plate had 4 sterility controls, 5 growth controls, 10 different concentrations of antimicrobial A alone, 7 different concentrations of antimicrobial B alone, and 70 different combinations of both antimicrobials A and B combined. Checkerboard plates performed in biologically-independent triplicates and were incubated overnight at 37°C. The OD584 values of each well were measured using a BMG Labtech FLUOstar Optima microplate reader.

### Analysis

After normalisation, wells that demonstrated an OD584 increase of ≥ 0.1 were considered positive for bacterial growth. Classification of the interaction of any two antimicrobials is based on the fractional inhibitory concentration (FIC):

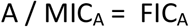

Where A is equal to the MIC of compound A when in combination and MIC_A_ is equal to the MIC of compound A when alone.

FIC Index (FICI) values were calculated as the FIC_A_ + FIC_B_ both from the same well. FICI values were deduced for all non-turbid wells along the turbidity/non-turbidity interface, as described by Bonapace *et al*., 2002. The lowest FICI value was used to characterise the interaction between the two antimicrobial compounds. FICI values were interpreted as synergistic if FICI ≤ 0.5, additive if 0.5 < FICI ≤ 1.0, indifferent if 1.0 < FICI ≤ 4 and antagonistic if FICI > 4.0. These commonly-used thresholds were selected to maintain comparability with other academic publications (13). Thresholds for additivity were included as non-selective, broad activity disinfectants were being tested.

### Time-Kill Assay

For further validation, disinfectant combinations that were identified as synergistic via the checkerboard method were tested for synergy via time-kill assays as described by CLSI (36). MHB cultures containing 5 × 10^5^ CFU/ml bacteria were exposed to either both antimicrobial compounds, one of the compounds alone or neither as a growth control. Antimicrobial concentrations were equal to those present in the well containing the lowest FICI value in the checkerboards previously conducted. Cultures had a final volume of 20 ml, with MHB used as the culture medium. Aliquots were taken at 0, 1, 3, 6, 12 and 24 hour time-points and the number of CFUs were quantified via culture analysis. All test conditions were conducted in triplicate.

A synergistic interaction was characterised as demonstrating a ≥2 log^10^ reduction in CFU/ml between the combination and its most active constituent alone after 24 hours. In addition, the number of CFU/ml must demonstrate a decrease of ≥ 2 log^10^ CFU/ml below the starting inoculum when exposed to the antimicrobial combination.

## ACKNOWLEDGEMENTS

This work was carried out by D. J. Noel as part of his Southampton NIHR Academy Fellowship to fulfil PhD degree training funded by NIHR Southampton Biomedical Research Centre and JVS Products Ltd who were not involved in the work nor in the writing of the paper.

